# Multi-parameter formulation development for an HIV-vaccine protein with direct validation of epitope binding integrity and stoichiometry

**DOI:** 10.1101/2020.11.30.403873

**Authors:** Corinna Popp, Philipp Schramm, Ralf Wagner, David Peterhoff, Christopher Battle, Christian Kleusch, Maximilian G. Plach

## Abstract

Vaccines are on the front line-of-defense against infectious diseases, ranging from threats we are familiar with, including polio, tuberculosis, or HIV, to novel and emerging threats such as SARS-CoV-2. Successful development of new protein-based vaccines requires sophisticated and efficient development of storage and formulation conditions. Here we demonstrate the combined power of 2bind’s sophisticated buffer matrix FORMOscreen^®^ and NanoTemper Technologies’ novel Prometheus Panta high-throughput Dynamic Light Scattering/Nano Differential Scanning Fluorimetry instrument. We show that this combination can comprehensively improve critical biophysical parameters for the HIV-1 vaccine BG505-SOSIP and find the optimal formulation condition with unmatched efficiency.

## Introduction

Vaccines are among the greatest biomedical achievements of the 20^th^ century, improving quality of life for billions around the world. Due to standard vaccination protocols, in the United States of America alone, the death toll from nine common infectious diseases (e.g. smallpox, diphteria, and measles) dropped from >1.1 million per year in the 1900s to just a bit over 7000 per year at the turn of the 21st century [1]. As of 2020, 97 FDA-licensed vaccines are available, targeting all sorts of bacterial and viral pathogens [2]. However, the need for the development of new and safe vaccines is higher than ever, as many threats are still looming out there: challenging but known pathogens like *Plasmodium* species, emerging pathogens like the *Severe Acute Respiratory Syndrome Coronavirus 2* (SARS-CoV-2), like *human immunodeficiency virus type-1* (HIV-1) [2, 3].

### Vaccine development and formulation

One of the key steps in vaccine development is the generation of an efficacious vaccine that is also stable enough to be stored and administered all over the world. To do so, successful vaccine formulation strategies must address many interrelated topics such as antigen stabilization, selection of appropriate adjuvants as well as the development of analytical methods for testing antigen stability and interaction with immune system components [4]. Ideally these topics are tackled in the initial stages of vaccine development in order to reduce late stage development attrition and failure rates.

### Prometheus Panta

A vaccine must be able to withstand the range of stresses it will encounter during production and long-term storage. Accelerated stability testing and biophysical characterization of candidate molecules is crucial to identifying and sorting out those prone to failure as early as possible. Many of the central questions in formulation development - thermal- and colloidal stability, solution homogeneity, aggregation propensity - can be addressed on a single instrument platform. Prometheus Panta automatically reports a thorough profile of the candidate molecule’s stability with a complete set of parameters for thermal unfolding, particle sizing, and self-interaction analysis. The instrument uses nanoDSF, DLS, and back-reflection technologies to provide results on thermal unfolding (melting temperature *T_m_*; onset of unfolding *T_onset_*; onset of heavy aggregation *T_turbidity_;* temperature at which average particle size begins to increase *T_size_*; reversibility of unfolding), isothermal particle analysis (hydrodynamic radius *r_h_*; polydispersity index *PDI*;), and self-interaction analysis (diffusion interaction parameter *k_D_*).

### Formulation development for an HIV-1 vaccine

A promising HIV-1 vaccine candidate is the HIV-1 envelope (*Env*) protein BG505 SOSIP.664 (BG505-SOSIP), which is currently being investigated in a phase-1 clinical trial [5]. BG505-SOSIP is a soluble, engineered version of the trimeric viral surface receptor gp140 and was developed to overcome failures of wild-type *Env* protein-vaccines to induce protective antibody responses [6]. In this white paper, we demonstrate the extensive capabilities of combining Prometheus Panta with the 2bind FORMOscreen^®^ buffer screen for finding the ideal formulation buffer for BG505-SOSIP. In this white paper, we show how to combine Prometheus Panta with the 2bind FORMOscreen^®^ to find a buffer that results in high thermal stability, a well-defined protein radius, excellent solution homogeneity and high stress-tolerance, while maintaining interaction with neutralizing anti-HIV-1 antibodies.

## Results and Discussion

### FORMOscreen^®^ buffer analysis

A key step in vaccine formulation development is finding a buffer composition that ensures optimal stability and activity of the protein vaccine and that serves as the basis for subsequent formulation refinement and storage testing. For this, combinatorial approaches are often performed in which many different buffer components are tested in varying combinations and subsequent experimental rounds. A similarly comprehensive but much quicker alternative is the 2bind FORMOscreen^®^ [7]: A collection of 96 ready-to-use buffers from patented formulations of developed antibodies that are approved by the U.S. Food and Drug Administration (FDA) or the European Medicines Agency (EMA).

One of the most routinely assessed stability parameters of therapeutic proteins and vaccines is their thermal stability, which can easily be monitored with thermal unfolding experiments. Higher thermal unfolding temperatures (*T_m_*) usually indicate a favorable stability equilibrium at ambient or storage temperatures. A nano-scale differential fluorimetry (nanoDSF) unfolding analysis of BG505-SOSIP in the 96 FORMOscreen^®^ buffers (Figure 1) revealed a large *T_m_* variation of 17.3°C. Some buffers enhanced BG505-SOSIP *T_m_* by up to +6.8°C (Table 1, compared to the base *T_m_* of 65.7°C in the 1 x PBS, pH 7.4 stock buffer; see Table S1 for full buffer compositions).

**Figure 1.**
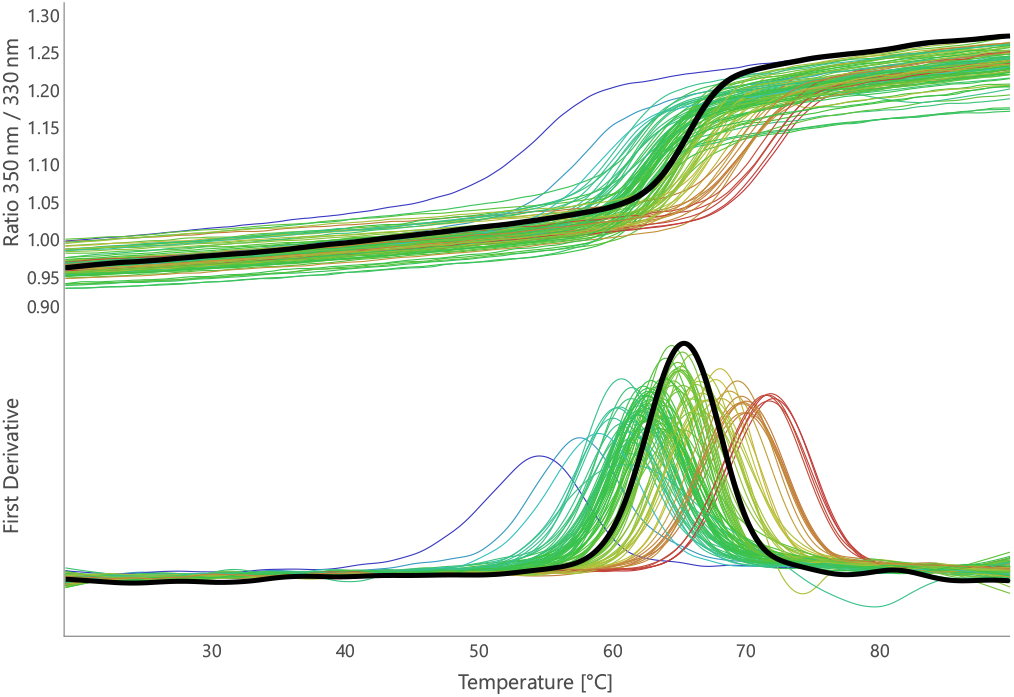
Initial nanoDSF FORMOscreen^®^ analysis of BG505-SOSIP. The fluorescence ratio signal and first derivative curve for the stock buffer are shown in black (1 x PBS, pH 7.4). The curves for the 96 FORMOscreen^®^ buffers are colored with respect to the resulting BG505-SOSIP melting temperature from blue (low) to red (high).

**Table 1.** DLS parameters of BG505-SOSIP in 20 selected buffers from the 2bind FORMOscreen^®^.

| FORMOscreen®<br>buffer number | $T_m$ /°C<sup>a</sup><br>(from [7]) | Hydrodynamic<br>radius /nm | PDI<sup>b</sup> |
| --- | --- | --- | --- |
| **10 buffers with highest BG505-SOSIP thermal stability ($T_m$)** |  |  |  |
| 9 | 72.50 ± 0.02 | 9.45 ± 4.70 | 0.28 ± 0.05 |
| 2 | 72.33 ± 0.18 | 7.62 ± 0.79 | 0.14 ± 0.04 |
| 35 | 72.06 ± 0.04 | 8.03 ± 1.36 | 0.27 ± 0.07 |
| 61 | 70.53 ± 0.03 | 6.89 ± 0.33 | 0.13 ± 0.04 |
| 36 | 70.45 ± 0.01 | 7.55 ± 0.27 | 0.09 ± 0.01 |
| 57 | 70.25 ± 0.03 | 7.18 ± 0.23 | 0.09 ± 0.02 |
| 49 | 70.20 ± 0.01 | 7.12 ± 0.43 | 0.11 ± 0.03 |
| 55 | 70.18 ± 0.15 | 6.61 ± 0.35 | 0.10 ± 0.05 |
| 7 | 70.06 ± 0.15 | 7.32 ± 0.88 | 0.26 ± 0.28 |
| 94 | 68.48 ± 0.01 | 8.78 ± 0.77 | 0.15 ± 0.03 |
| **10 buffers with lowest BG505-SOSIP thermal stability ($T_m$)** |  |  |  |
| 46 | 61.94 ± 0.03 | 7.08 ± 0.28 | 0.10 ± 0.03 |
| 75 | 61.69 ± 0.07 | 8.66 ± 2.15 | 0.21 ± 0.05 |
| 45 | 61.29 ± 0.23 | 10.5 ± 6.32 | 0.36 ± 0.04 |
| 32 | 61.20 ± 0.07 | 5.85 ± 0.30 | 0.10 ± 0.02 |
| 48 | 61.06 ± 0.02 | 8.21 ± 0.43 | 0.10 ± 0.04 |
| 70 | 60.83 ± 0.08 | 12.0 ± 9.63 | 0.47 ± 0.07 |
| 71 | 60.59 ± 0.04 | 7.19 ± 0.33 | 0.41 ± 0.07 |
| 34 | 59.46 ± 0.31 | 6.64 ± 0.16 | 0.09 ± 0.01 |
| 73 | 58.23 ± 0.16 | 8.88 ± 3.42 | 0.26 ± 0.03 |
| 16 | 55.20 ± 0.03 | 7.18 ± 1.67 | 0.10 ± 0.01 |

<sup>a</sup> Thermal melting temperature
<sup>b</sup> polydispersity index

Thus, in a single experiment, the top candidate buffers (Table 1, top 10) can be differentiated from the buffers that result in sub-optimal vaccine stability (Table 1, bottom 10). Importantly, the full FORMOscreen^®^ analysis of BG5050-SOSIP (duplicate experiments) required only 80 *μg* protein and was completed within a single day (Table 2).

**Table 2.** Sample and time requirements for FORMOscreen^®^/Prometheus Panta buffer optimization.

| Experiment | Parameters<sup>a</sup> | Sample requirement | Time requirement |
| --- | --- | --- | --- |
| Initial 96-condition FORMOScreen® nanoDSF analysis | $T_m$, $T_{turbidity}$ | 80 μg | 1 working day |
| 20-condition FORMOScreen® DLS refinement | $r_h$, PDI | 60 μg | 1 working day |
| 1-condition accelerated-stress matrix analysis | $r_h$, PDI | 280 μg | 2 working days (excluding storage time) |
| Continuous variation binding analysis | $r_h$, Stoichiometry | 10 μg | <1 working day |

<sup>a</sup> $T_m$ : Thermal melting temperature, $T_{turbidity}$ : macroscopic aggregation temperature, $r_h$ : hydrodynamic radius, PDI: polydispersity index.

### DLS-characterization of BG505-SOSIP

The ten buffers with the highest and lowest BG505-SOSIP *T_m_* were selected for Prometheus Panta Dynamic Light Scattering (DLS) characterization with respect to BG5050-SOSIP hydrodynamic radius and polydispersity index (PDI) (Table 1). Interestingly, no clear correlations were observed between *T_m_* on the one hand and the hydrodynamic radius and PDI on the other hand. This suggests that while *T_m_* is a suitable parameter for judging the thermal stability of a protein and, to a first approximation, the equilibrium between folded and unfolded state at lower temperatures, it should not be taken as a measure of protein homogeneity at ambient temperatures.

Two buffers illustrate this observation very well: Buffer 9 resulted in the highest overall *T_m_* of BG505-SOSIP (Table 1). However, the hydrodynamic radius of BG505-SOSIP in buffer 9 was also among the largest radii observed, second highest standard deviation and the PDI of 0.28 indicated a polydisperse sample quality. Figure 2 shows the DLS and nanoDSF data in detail. The relative frequency plot makes clear that in addition to the main BG505-SOSIP species with *r_h_* of 7-8 nm, additional, larger species (*r_h_* around 100 nm) are present. These larger species lead to a higher mean cumulant *r_H_* and a higher PDI, as the sample is present in a more heterogenous state. The very high sensitivity of Prometheus Panta even allows for the detection of sugar species in the sample. Buffer 9 contains 80 mg/ml sucrose (Table S1) and the sugar shows up as an additional peak with *r_h_* <1 nm. Small *r_h_* values <1.5 nm were not included in the calculation of the mean *r_h_* and PDI, but are shown in the size distribution analysis.

**Figure 2.**
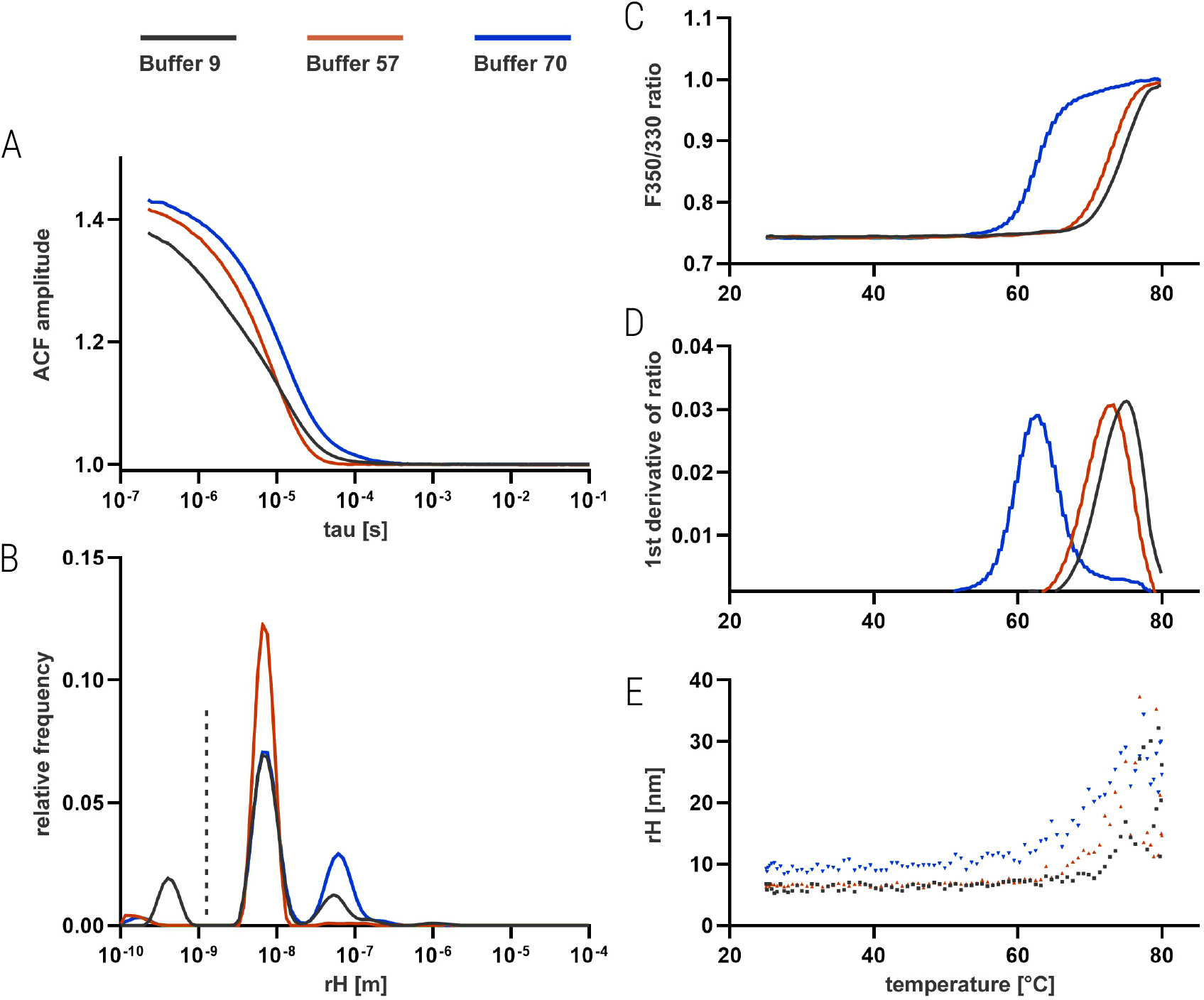
Exemplary DLS-based isothermal particle characterization and combined DLS/nanoDSF stability characterization of BG505-SOSIP in selected FORMOscreen^®^ buffers. **(A)** Autocorrelation functions of isothermal DLS analysis of BG505-SOSIP in buffers 9, 57, and 70. **(B)** Size distribution fits with relative frequency of detected particle sizes plotted against the mean hydrodynamic radius from cumulant analysis. The dashed vertical line visualizes the 1.5 nm cut-off for the cumulant fit. **(C)** F350/F330 ratio curves from the thermal unfolding experiment of BG505-SOSIP. **(D)** First derivative curves of the thermal unfolding F350/F330 data. **(E)** Temperature-dependent variation of hydrodynamic radius.

In contrast to buffer 9, buffer 57 resulted in a 2.25°C lower melting temperature but excellent DLS parameters (Table 1). The hydrodynamic radius of BG505-SOSIP was very well defined at 7.18 nm and the PDI of only 0.09 indicated a very homogeneous solution. Consequently, the DLS data (Figure 2) only show one well defined protein species in the relative frequency plot of the size distribution analysis.

To complete the picture, the detailed DLS analysis confirmed that in some cases there is a clear correlation between low melting temperature and sub-optimal solution properties. For example, buffer 70 resulted in a very large cumulant protein radius (12.0 nm, Table 1), which is also evident from the relative frequency plot (Figure 2) and also significantly destabilized BG505-SOSIP.

A unique feature of Prometheus Panta is the ability to collect all revelant DLS data (*r_h_* and PDI) not only isothermally, but across a temperature gradient between 25 and 95°C. In general, the temperature at which the average particle size begins to increase (*T_size_*) correlates well with the onset of thermal unfolding (*T_onset_*). For example, *T_size_* is higher for buffers 9 and 57 than for buffer 70, which also had the lowest *T_m_* of those three buffers. Another significant advantage over other DLS instruments is the ultra-low sample consumption of Prometheus Panta. The full DLS-characterization of BG505-SOSIP in the 20 selected FORMOscreen^®^ buffers was done with just 60 *μ*g of protein and within a 2 hour measurement time.

### Accelerated-stress matrix analysis

Up to this point, a promising candidate for a BG505-SOSIP formulation buffer has been identified in FORMO-screen^®^ buffer 57. But for a successful application of buffer 57, more knowledge about long-term effects on BG505-SOSIP are required. For this, accelerated-stress studies are usually done to test the stress-response of the vaccine candidate. As there are no clearly defined regulatory guidelines for accelerated-stress analysis of vaccine proteins, we selected certain prevalent testing conditions from the literature [8]: (i) Thermal stress at 25, 40, and 60°C for up to 14 days, (ii) oxidative stress in presence of hydrogen peroxide and 25, 40, and 60°C for up to 14 days, (iii) mechanical stress by shaking and vortexing, and (iv) freeze-thaw stress (Figure 3).

**Figure 3.**
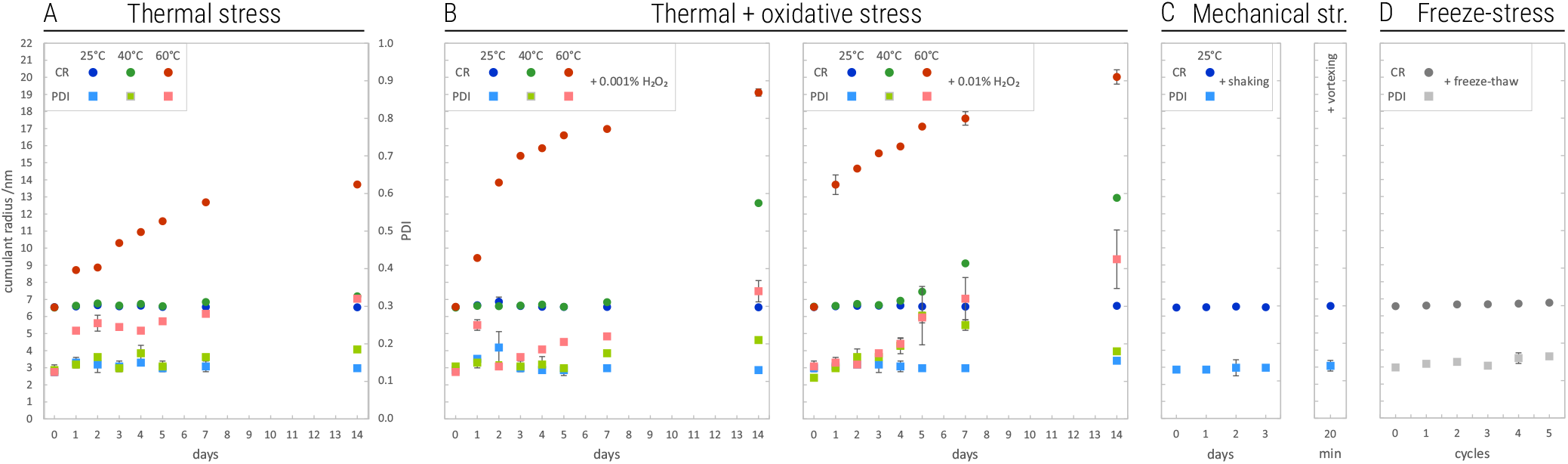
Accelerated-stress analysis of BG505-SOSIP. **(A)** Thermal stress at 25, 40, and 60°C. **(B)** Combined thermal and oxidative stress at 25, 40, and 60°C in presence of 0.001% or 0.01% hydrogen peroxide, respectively. **(C)** Mechanical stress at 25°C by shaking at 350 rpm or vortexing. **(D)** 5-cycle freeze-thaw stress (flash-freezing in liquid nitrogen, storage at −80°C, thawing to room temperature). Vertical axes for panels B-D as in panel A. Data are mean ± SD from at least two independent experiments. The majority of cumulant radius- and PDI-SD are <0.05 nm and <0.01, respectively.

The FORMOscreen^®^ buffer 57 is very well suited for long-term storage of BG505-SOSIP. At 25°C and 40°C storage temperature, the hydrodynamic radius of BG505-SOSIP and the solution PDI do not significantly change over the course of two weeks (Figure 3A). Only after prolonged storage at 60°C the hydrodynamic radius roughly doubles, which indicates conformational changes or partial protein unfolding and the PDI increases to >0.3, indicating an inhomogeneous solution quality. Even under oxidative stress (Figure 3B,C), radius and PDI are not affected at all at 25°C and only slightly affected at 40°C. The observed radius increase at 60°C is more pronounced under additional oxidative stress. Buffer 57 also enables BG505-SOSIP to tolerate severe mechanical stress at ambient temperature (Figure 3D,E; vortexing for 20 minutes) and five freeze-thaw cycles; both hydrodynamic radius and PDI are practically identical to their pre-stress values.

Importantly, the full accelerated-stress study of BG505-SOSIP in buffer 57 could be done with only 280 *μ*g of protein due to the exceptional sensitivity and ultra-low samples consumption of Prometheus Panta. Moreover, not considering incubation and storage times, the analysis took only two working days Table 2.

### DLS-based interaction analysis of BG505-SOSIP - antibody complexes

A successful vaccine formulation must also ensure induction of proper immune response. A critical quality control criterion in modern reverse vaccinology is high affinity binding of the vaccine to well characterized protecting antibodies. While many different assay formats and technologies can be used to test vaccine-antibody binding, the same instrument platform, herein used for optimizing formulation conditions, can also be applied for direct validation of BG505-SOSIP epitope binding integrity. This rather uncommon approach does not require any labeling, immobilization or any other treatment of vaccine or antibody. Moreover, this method can be performed directly on the same Prometheus Panta instrument that was used for formulation development.

The method of choice is “continuous variation” [9], in which the total molar concentration of both interaction partners (BG505-SOSIP and neutralizing IgG1*κ* antibodies) is kept constant but the molar ratio between them is continuously varied from 100% BG505-SOSIP to 100% antibody over a large number of samples (typically >20). For each sample, the cumulant *r_h_* was determined via Prometheus Panta DLS size analysis and plotted against the molar ratio of antibody:total protein (Figure 4).

**Figure 4.**
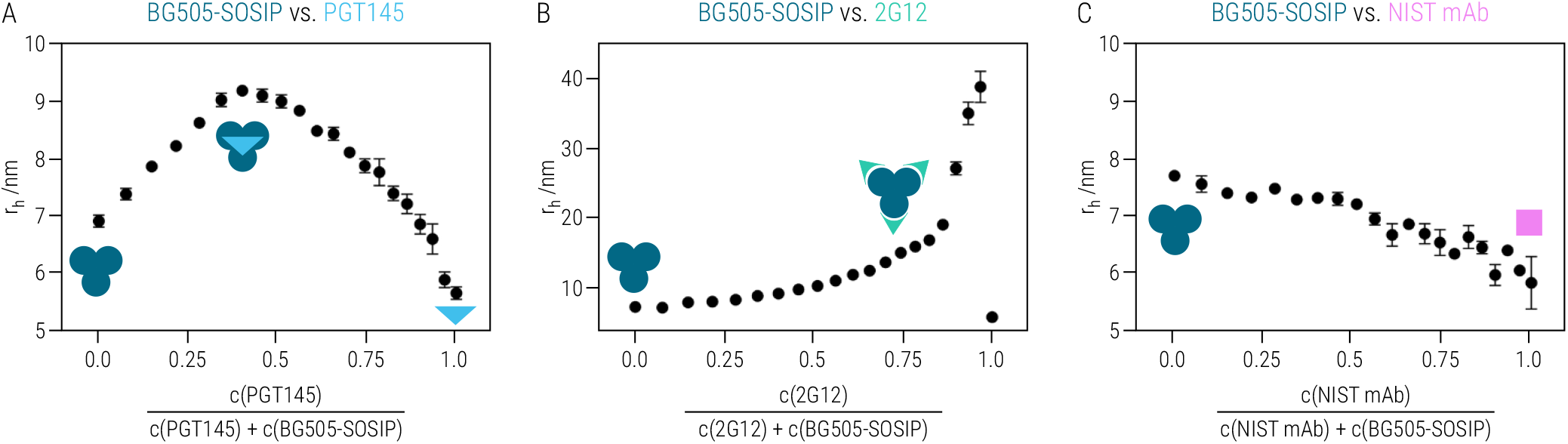
Continuous variation binding experiments. The molar ratio of BG505-SOSIP and neutralizing antibodies (**A**, **B**) and a control antibody, respectively (**C**), was varied between 0 and 1 and the cumulant hydrodynamic radius for 21 different molar ratios was determined.

First, we tested binding of the broadly-neutralizing antibody PGT145, which is known to bind exclusively to BG505-SOSIP trimers [10]. The continuous variation plot (Figure 4A) has a maximum at a molar ratio of 0.5, indicating a 1:1 interaction stoichiometry between the BG505-SOSIP trimers and PGT145. The average hydrodynamic radius at this molar ratio was 9.0±0.1 nm. The samples with only BG505-SOSIP or only PGT145 antibody resulted in the expected radii; e.g. 7.0±0.1 nm for BG505-SOSIP and 5.7±0.1 nm for the antibody.

Next, a different antibody, 2G12, was tested, which binds each BG505-SOSIP protomer individually via an epitope on the outer domain of the gp120 subunits [10]. As expected, the average radius increases with increasing molar ratio (Figure 4B) and seems to approach a maximum around a molar ratio of 0.75, congruent with the described 1:3 binding stoichiometry. Above the molar ratio of 0.75, however, the average radius strongly increases to around 40 nm. A possible explanation for this observation is the ability of 2G12 to bind BG505-SOSIP with a second, much weaker affinity interaction. This may lead to inter-molecular cross-linking of two or more BG505-SOSIP trimers by 2G12. Even at low, absolute concentrations of the 2G12 (i.e. high molar ratios), this behavior was observed. This is not surprising, because larger particles scatter light much more intensely (according to Rayleigh approximation, the scattering intensity is proportional to d^6^) [11] and thus just a few cross-linked, large protein-antibody complexes can mask all other scattering signals in the average radius analysis. Consequently, without any BG505-SOSIP, the cumulant *r_h_* for 2G12 drops to 5.76 nm. This showcases one limitation of such a continuous variation binding assay: The hydrodynamic radii of the involved species must not deviate from each other with a factor >2 (or in other words, the molecular weights of the involved species should not be more different than 5-fold) [9]. A control with a non-related antibody (The NIST monoclonal antibody (NISTmAb) reference material, RM 8671) [12] did not show signs of an interaction and the average hydrodynamic radius continuously changed from the BG505-SOSIP radius to that of the antibody (Figure 4C).

## Conclusions

For improving the thermal stability, solution homogeneity, stress-tolerance, aggregation propensity and epitope accessibility of HIV spike-like vaccine candidate BG505-SOSIP, we tested the sophisticated buffer matrix FORMOscreen^®^ on a Prometheus Panta instrument. FORMOscreen^®^ buffer 57 has been shown comprehensively to be a perfect candidate for formulating the BG505-SOSIP HIV-1 vaccine candidate: The protein is highly stable in this buffer, has a very well-defined hydrodynamic radius, is very homogeneous, and keeps these properties even under stress. The results of this showcase study impressively demonstrate the accelerated process of finding optimal formulation conditions at unmatched efficiency. We successfully improved critical biophysical parameters of the antiviral vaccine candidate with minimal sample consumption and extremely shortened experimental measurement time. Ultimately, optimal formulation conditions improve the likeliness of an antiviral vaccine candidate to be successfully developed and ensure that its beneficial, protective effect can be safely delivered to patients.

## Materials and Methods

### Thermal unfolding analysis

Thermal unfolding of BG505-SOSIP was determined with a heating ramp of 1°C/min and 20% sensitivity setting using high-sensitivity capillaries (NanoTemper Technologies, Munich, Germany; Cat# PR-C006). BG505-SOSIP was diluted to 700 nM in selected FORMOscreen^®^ buffers (2bind, Regensburg, Germany, Cat# 2BBT-001).

### Dynamic Light Scattering analysis

BG505-SOSIP hydrodynamic radius and PDI in FORMOscreen^®^ buffers (2bind, Regensburg, Germany, Cat 2BBT-001) were determined with DLS using Prometheus Panta’s Size Analysis function. Isothermal DLS scans (10 acquisitions, 5 s each, 20% LED intensity, 100% DLS-Laser intensity) were performed at 25°C or 80°C. Measurements at 80°C required capillary sealing (Capillary Sealing Paste and Capillary Sealing Applicators; NanoTemper Technologies, Munich, Germany; Cat# PR-P001, PR-P002). BG505-SOSIP was diluted to 700 nM in selected FORMOscreen^®^ buffers. All plasticware was rinsed with 0.22 micron-filtered mQ-H2O before sample preparation. Buffer viscosity base values were provided to the Prometheus Panta control software.

### Accelerated-stress assays

Isothermal DLS scans of BG505-SOSIP in sterile-filtered FORMOscreen^®^ buffer 57 (2bind, Regensburg, Germany, Cat# 2BBT-001) were performed on Prometheus Panta as described above. BG505-SOSIP was diluted to 700 nM, centrifuged (30 min at 4°C, 18000 rcf), and treated with the following conditions before DLS analysis. (A) Thermal stress: Incubation at 25°C, 40°C, and 60°C for 0-14 days as well as incubation at 95°C for 2 h and 24 h. (B) Oxidative stress: Incubation at 25°C, 40°C, and 60°C for 0-14 days in presence of 0.001% or 0.01% hydrogen peroxide. (C) Mechanical stress: Incubation at 25°C with shaking at 350 rpm for 0-72h as well as vortexing for 20 min. (D) Freeze/thaw stress: Snap-freezing in liquid nitrogen followed by storage at −20°C for at least 1 h and thawing to room temperature.

### DLS-based interaction analysis

Interaction between BG505-SOSIP versus monoclonal IgG1*κ* antibodies (2G12, PGT145 and NIST mAb RM8671) was tested in a concentration gradient experiment, using Prometheus Panta’s isothermal, high sensitivity DLS scan mode. In short, concentrations of the two interaction partners were continuously varied in a concentration gradient against each other. The interaction experiment was prepared in 384-well, non-binding surface, assay plates (Corning Inc., Kennebunk ME, USA, product # 4513). Based on the nanoDSF and DLS FORMOscreen^®^ results, buffer 57 was selected to test the interaction of different IgG1*κ* antibodies vs BG505-SOSIP. Stock solutions of all interaction partners were diluted in buffer 57 to final concentrations for BG505-SOSIP of 447 nM, and 700 nM for the monoclonal antibodies. Each well was filled to a total volume of 30 *μ*l, with well 1 containing BG505-SOSIP only and well 21 containing only monoclonal antibody. All wells in between contained decreasing BG505-SOSIP and increasing antibody concentrations (e.g. well 2 424.65 nM BG505-SOSIP vs 35 nM antibody, well 3 402.3 nM BG505-SOSIP vs 70 nM antibody, well 20 22.35 nM BG505-SOSIP and 665 nM antibody, and so on). Each concentration gradient experiment was incubated for 30 minutes at room temperature, samples filled into Prometheus High Sensitivity capillaries and transferred to Prometheus Panta instrument. High sensitivity DLS scans were performed in duplicates at 25°C, viscosity of buffer 57 was taken into account (see also Dynamic Light Scattering Analysis), and autocorrelation functions were fitted using cumulant analysis method. Obtained apparent hydrodynamic radii were plotted as a function of the molar ratio between BG505-SOSIP and respective antibody.

## Supporting Information

**Table S1.** 2bind FORMOscreen^®^ buffer compositions.

| Buffer number | Buffer composition |
| --- | --- |
| 9 | 0.4 mg/ml Polysorbate 80, 80 mg/ml Sucrose, 2.7 mg/ml Sodium citrate dihydrate, 0.16 mg/ml Citric acid monohydrate, pH 6.5 |
| 2 | 0.2 mg/ml Polysorbate 80, 70 mg/ml Trehalose dihydrat, 5.6 mg/ml Sodium citrate dihydrate, 0.21 mg/ml Citric acid monohydrate, pH 6.6 |
| 35 | 0.1 mg/ml Polysorbate 80, 62 mg/ml Sucrose, 4.91 mg/ml Sodium citrate dihydrate, 0.2 mg/ml Citric acid monohydrate, pH 6.6 |
| 61 | 0.2 mg/ml Polysorbate 80, 22.5 mg/ml Glycine, 0.44 mg/ml Citric acid monohydrate, pH 5.7 |
| 36 | 10 mg/ml Sucrose, 1.5 mg/ml Sodium chloride, 5.75 mg/ml Lysine hydrochloride, 13.52 mg/ml Sodium citrate dihydrate, 0.86 mg/ml Citric acid monohydrate, pH 6.3 |
| 57 | 0.1 mg/ml Polysorbate 80, 2.34 mg/ml Sodium chloride, 9.11 mg/ml Mannitol, 9.98 mg/ml Glycine, 2.55 mg/ml Sodium citrate dihydrate, 0.28 mg/ml Citric acid monohydrate, pH 6.0 |
| 49 | 0.34 mg/ml Polysorbate 20, 60 mg/ml Trehalose dihydrat, 6.56 mg/ml Sodium phosphate monobasic dihydrate, 1.2 mg/ml Sodium phosphate dibasic anhydrous, pH 6.1 |
| 55 | 0.2 mg/ml Polysorbate 80, 2.92 mg/ml Sodium chloride, 30 mg/ml Mannitol, 0.01 mg/ml Pentetic acid, 5.88 mg/ml Sodium citrate dihydrate, pH 6.0 |
| 7 | 0.4 mg/ml Polysorbate 20, 60 mg/ml Trehalose dihydrat, 6.56 mg/ml Sodium phosphate monobasic dihydrate, 1.2 mg/ml Sodium phosphate dibasic anhydrous, pH 6.2 |
| 94 | 47.62 mg/ml Sucrose, 0.55 mg/ml Sodium chloride, 3.29 mg/ml Sodium phosphate monobasic dihydrate, pH 7.4 |
| 46 | 0.2 mg/ml Polysorbate 20, 0.3 mg/ml L-Histidine, 1.7 mg/ml Histidine Hydrochlorid monohydrate, 2.9 mg/ml Sodium chloride, 13.7 mg/ml Mannitol, 7.5 mg/ml Glycine, pH 5.4 |
| 75 | 0.15 mg/ml Polysorbate 80, 0.28 mg/ml L-Histidine, 1.6 mg/ml Histidine Hydrochlorid monohydrate, 45 mg/ml Sorbitol, pH 5.5 |
| 45 | 0.2 mg/ml Polysorbate 80, 70 mg/ml Sucrose, 1.55 mg/ml L-Histidine, pH 5.5 |
| 32 | 3.1 mg/ml L-Histidine, 26.1 mg/ml L-Arginine, 26.1 mg/ml L-Arginine hydrochloride, 0.5 mg/ml Poloxamer 188, pH 6.0 |
| 48 | 0.1 mg/ml Polysorbate 20, 10 mg/ml Histidine Hydrochlorid monohydrate, 100 mg/ml Trehalose dihydrat, pH 5.4 |
| 70 | 0.1 mg/ml Polysorbate 80, 25.43 mg/ml Proline, 1.2 mg/ml Sodium acetate, pH 5.0 |
| 71 | 0.1 mg/ml Polysorbate 80, 25 mg/ml Proline, 1.2 mg/ml Sodium acetate, pH 5.0 |
| 34 | 1 mg/ml Polysorbate 80, 6.15 mg/ml Sodium chloride, 12 mg/ml Mannitol, 0.3 mg/ml Sodium citrate dihydrate, 1.3 mg/ml Citric acid monohydrate, 0.85 mg/ml Sodium phosphate monobasic dihydrate, 1.24 mg/ml Sodium phosphate dibasic anhydrous, pH 4.7 |
| 73 | 0.1 mg/ml Polysorbate 20, 24 mg/ml Proline, 4.33 mg/ml Glutamate, pH 4.8 |
| 16 | 7.31 mg/ml Sodium chloride, 1.36 mg/ml Sodium acetate, pH 4.6 |

